# Quantitative sequence basis for the *E. coli* transcriptional regulatory network

**DOI:** 10.1101/2022.02.20.481200

**Authors:** Sizhe Qiu, Cameron Lamoureux, Amir Akbari, Bernhard O. Palsson, Daniel C. Zielinski

## Abstract

The transcriptional regulatory network (TRN) of *E. coli* consists of thousands of interactions between regulators and DNA sequences. Inherently the DNA sequence is the primary determinant of the TRN; however, it is well established that the presence of a DNA binding motif does not guarantee a functional regulatory protein binding site. Thus, the extent to which the TRN architecture can be predicted by the genome DNA sequence alone remains unclear. Here, we developed machine learning models that predict the TRN structure of *E. coli* based on genome sequence. Models were constructed successfully (cross-validation AUROC >= 0.8) for 84% (57/68) of valid *E. coli* regulons identified from top-down analysis of RNA-seq data. We found that: 1) While regulatory motif strength is the most important sequence feature for determining regulon membership, additional features such as DNA shape substantially influence membership; 2) complex regulons involving multiple interacting regulators could be unraveled by machine learning; 3) investigating regulons where initial ML models failed revealed new regulator-specific sequence features that improved model accuracy. Finally, while regulon structure can appear to be variable across estimation methods and strains, we found that strong regulatory sequence features underlie both the genes that appear most consistently in regulons across estimation methods as well as the core regulon genes in the Fur pan-regulon. This work develops a quantitative understanding of the sequence basis of the TRN and suggests a path towards computationally-guided control of transcriptional regulation for synthetic biology applications.

## Introduction

Microbial gene expression is tightly controlled to maintain fitness under diverse conditions. Understanding these regulatory mechanisms is critical for enabling the modification of gene expression for synthetic biology applications. Among the processes through which expression is controlled, transcription initiation stands out as a critical and highly regulated process (1). In transcription initiation, the RNA polymerase recognizes the promoter sequence upstream of a coding region, with the polymerase sigma factor binding to a conserved motif followed by the formation of an open complex of the RNA polymerase and DNA sequence (1). Transcription factors (TFs) often bind to promoter regions at TF-specific motifs to activate or repress transcription by altering the recruitment of the polymerase complex to the promoter. Regulons, or sets of genes that are transcriptionally regulated by the same protein, have been identified for at least 115 TFs and 7 sigma factors in *E. coli* (2). This set of regulons constitutes the transcriptional regulatory network (TRN) of an organism, representing thousands of interactions between regulators and promoter sequences.

A variety of experimental and computational approaches have contributed to our current knowledge of the structure of the TRN of *E. coli.* Chromatin immunoprecipitation (ChIP) is a bottom-up method for TRN elucidation that identifies TF binding sites across the genome by crosslinking bound proteins to DNA and then sequencing the captured DNA segments (3). These binding sites are captured as “peaks” with a particular signal-to-noise (S/N) ratio, depending on the number of reads of a given sequence present after alignment. As ChIP experiments alone do not indicate whether the TF binding regulates expression of the gene, ChIP is often paired with differential expression analysis after TF knockout to validate that the TF binding is regulatory (4). A set of binding sites above a certain S/N enrichment threshold that also affect gene expression comprise a bottom-up, component-by-component estimation of the TF regulon. Over the years, this process has been repeated for many of the major *E. coli* transcription factors, enabling a bottom-up, ChIP-based estimation of the TRN of *E. coli (5, 6)*.

As an alternative approach, network inference methods have been used to provide topdown estimates of the TRN based on analysis of RNA-seq compendia (7). In particular, independent component analysis (ICA) of a large *E. coli* RNA-seq dataset has recently provided a highly informative estimation of the TRN of *E. coli (8)*. ICA is an unsupervised machine learning method similar to PCA that identifies sets of coordinated variables, termed components; however, while PCA maximizes variance in the data explained by components, ICA maximizes statistical independence between the components it calculates (9). ICA has been applied to large RNA-seq expression compendia to identify independently modulated groups of genes, termed iModulons, in multiple organisms (10). These iModulons tend to overlap significantly with ChIP-determined regulons, supporting the validity of ICA-calculated TRNs. To better clarify the analogy to ChIP-determined regulons, in this work we will refer to iModulons as ICA regulons.

While our understanding of the TRN of *E. coli* likely exceeds that of any other organism, there is still much that we do not understand regarding how the TRN is encoded in the genome itself (11). While we have identified binding sequence motifs for many transcription factors, the presence of a motif in the promoter sequence does not guarantee that a regulator will bind or influence gene expression(11). Thus, we still lack a quantitative understanding of how promoter sequence encodes a functioning regulatory site.

In this study, we developed a quantitative understanding of the *E. coli* TRN by constructing machine learning models that predict regulon membership based on promoter sequence. With these models, we identified important features that determine regulon membership. We utilized these machine learning models to understand the basis for complex top-down regulons involving multiple regulators. Further, we investigated differences between bottom-up and top-down TRNs to find the general factors underlying TRN estimates from these methods. Furthermore, we expanded the scope of our analysis to multiple strains of *E. coli*, and we explained variation of regulatory activity across strains using sequence features, enabling the prediction of TRN structure in other strains.

## Results

### Promoter sequence features can quantitatively predict a large part of the *E. coli* TRN

First we sought to determine the degree to which the *E. coli* TRN can be predicted based on promoter sequence by building machine learning models for ICA regulons. We developed a workflow to quantify the gene promoter sequence into a sequence feature matrix, and then trained a logistic regression (LR) classifier to predict ICA regulon membership (**Figure 1A-E**). The feature matrix contains: 1) sigma factor related features, including motif score and Hamming distance of −10/−35 boxes, spacer length, AT content, 2) TF related features, including motif score and shape features, and 3) genome organization features, including strand direction and binding site distance to transcription start site (**Figure 1B**).

**Figure 1.**
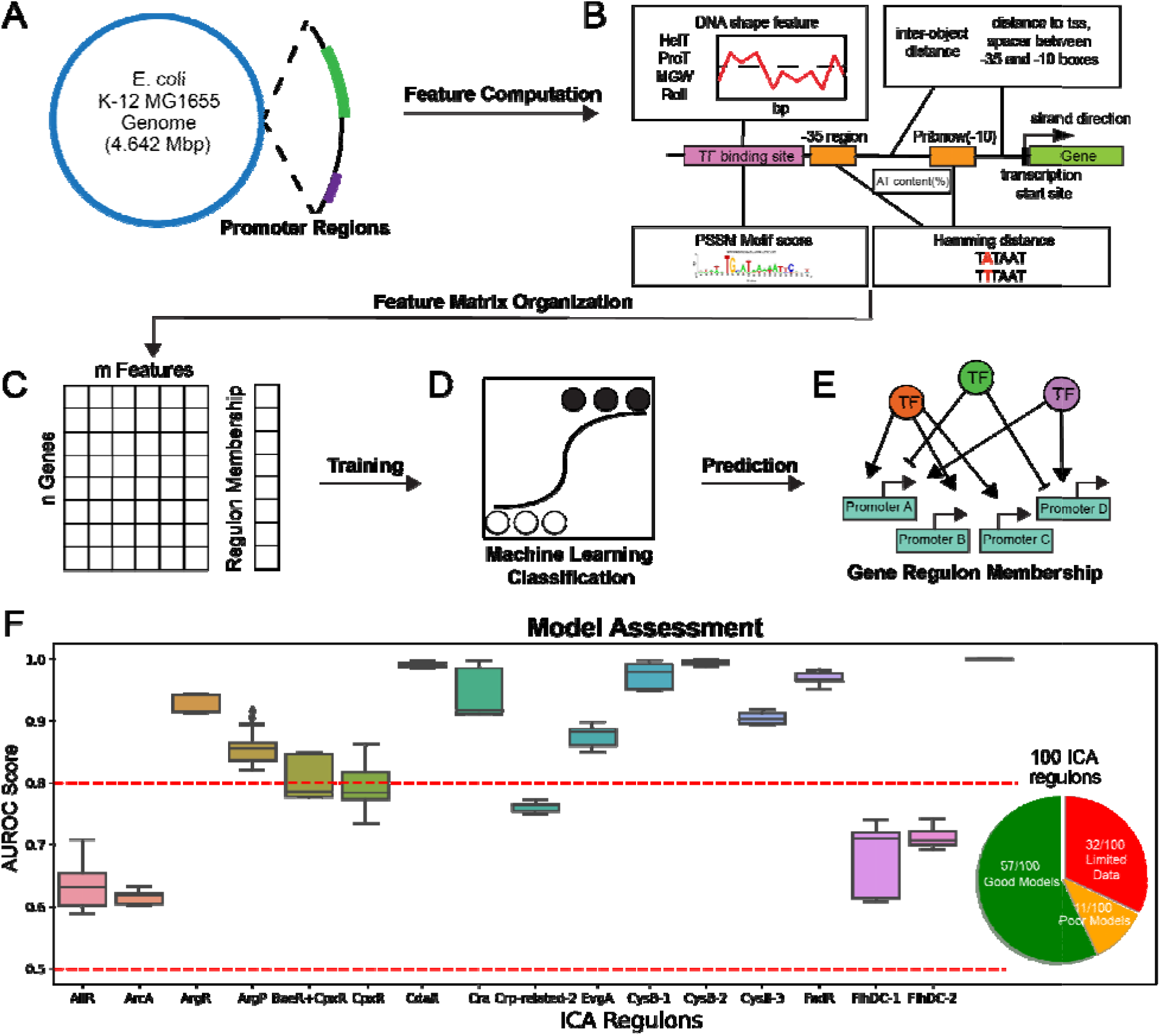
Sequence-based TRN prediction workflow and model assessment. (A) Gene promoter regions are extracted from the genome sequence of *E. coli* K12 MG1655. (B) Feature computation from the promoter sequence extracted from the genome. (C) Feature matrix organization and ICA regulon membership as the target label. (D) Logistic regression classifier trained on promoter features. (E) The predicted ICA TRN. (F) Model assessment by area under ROC curve and overall predictive model status. Cutoff = 0.8 for good models and 0.5 for baseline models.

The current version of the *E. coli* K12 MG1655 TRN estimated by ICA contains 100 total regulons (12). Models could not be constructed for 32 of these regulons due to size limitations or poor transcription start site annotation. Model assessment by 5-fold cross-validation showed that models for 57 of the remaining 68 regulons had AUROC scores exceeding a 0.8 threshold (**Figure 1F**), while models for 11 regulons performed under this threshold (**Figure 1F**). In short, more than 50% of the *E. coli* TRN could be predicted well by promoter sequence features using an LR classifier.

By examining the resulting machine learning models, we then determined the most essential sequence features for defining regulon membership. We discuss three representative examples of highly accurate models, including both TF and sigma factor-associated regulons (**Figure 2A-C**). SHAP values indicate that regulator binding site motif scores and shape features are important features (**Figure 2D-F**). Consistent with feature importance assessment, motif scores and shape features display distinct distributions for genes in the regulon (**Figure 2G-I**). With respect to sigma factor associated ICA regulons, distinctively high motif scores and low Hamming distances of −10/−35 boxes for RpoH contribute to the prediction of genes in RpoH ICA regulon, as both metrics give good characterization of sigma factor binding affinity (**Figure 2C**). In general, motif scores or Hamming distances of TF and sigma factors, and DNA shape features are essential to determine the ICA TRN.

**Figure 2.**
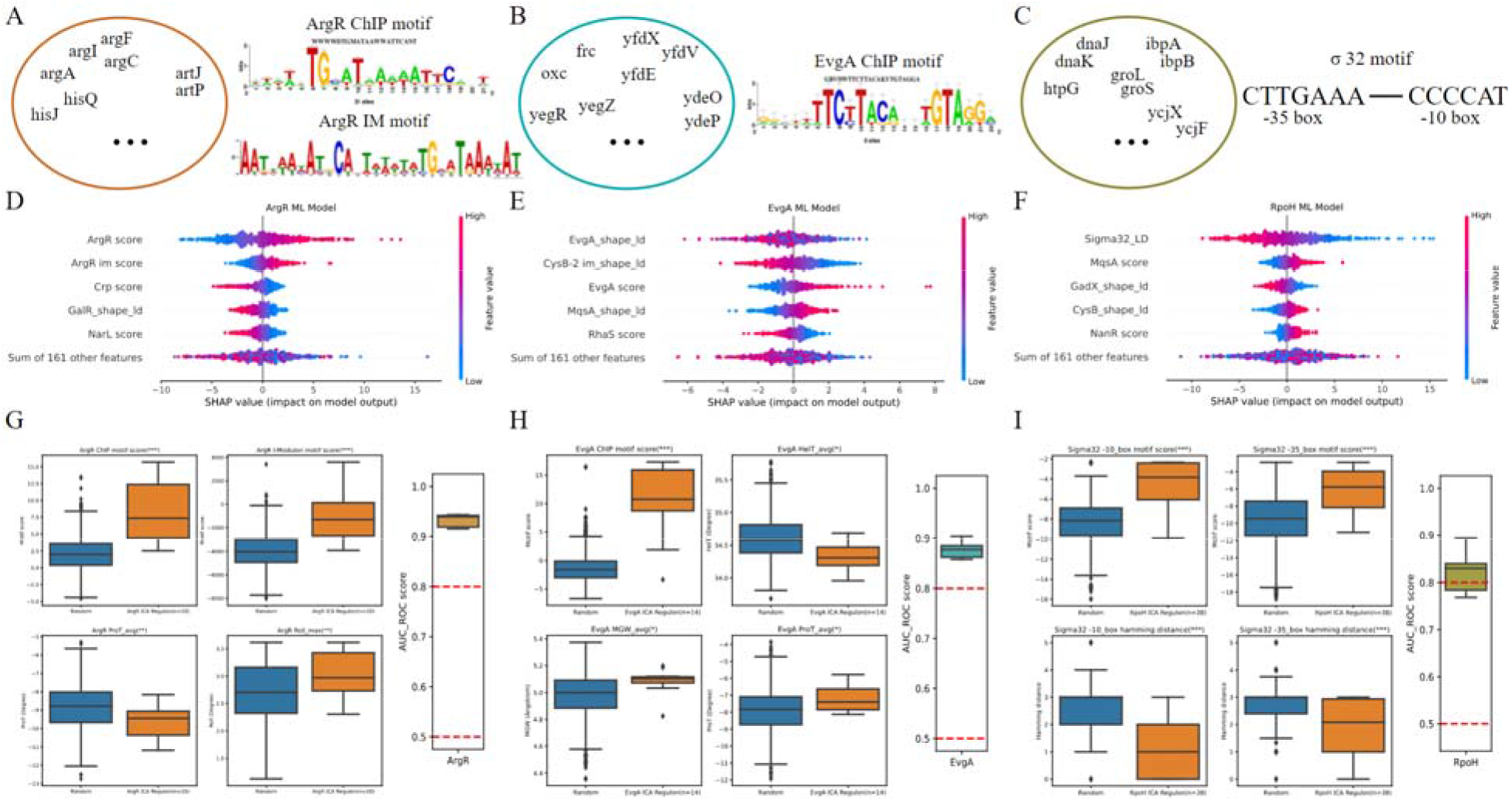
Examples of predictive models’ determining features: feature distributions, feature importances assessed by SHAP values, and model performances (A) ChIP and ICA regulon motifs of ArgR. (B) ChIP motif of EvgA. (C) Consensus sequences of RpoH’s −35/−10 boxes. (D) Feature importances for ArgR. (E) Feature importances for EvgA. (F) Feature importances for RpoH.(G) Feature distributions for ArgR. (H) Feature distributions for EvgA. (I) Feature distributions for RpoH.

### Underperforming models could be improved through additional TF-specific features

ArcA is an important global regulator involved in redox regulation(4). The initial machine learning model for ArcA showed relatively poor performance (**Figure 1**), indicating that features we used, which included the ArcA binding motif determined by ChIP, were not sufficient to determine the ICA regulon membership. In addition, we found that the overlap between ICA and ChIP-determined regulons was relatively low (**Figure 3B**), which could explain why the ChIP-determined motif was not useful in prediction for the ICA-determined ArcA regulon. A literature investigation suggested that the binding site of ArcA has great diversity, and involves a variable number of direct repeats (DRs) of the ArcA motif (13). Thus, rather than calculating a single ArcA motif score, it may be more biochemically accurate to define scores for multiple possible DR motifs with letter probability matrices (**Figure 3A**). Using position-specific scoring matrices (PSSMs) generated from letter probability matrices, we aligned DR motifs to promoter sequences (**Figure 3E**). Comparing predictive models generated using multiple DR motifs and those using only ChIP motifs separately, we found that ArcA model performance was greatly improved by the inclusion of DR motif scores (**Figure 3C**). The lower distributions of 3DR and 4DR motif scores in ArcA ChIP regulon suggest that it fails to capture genes with promoters containing extended DR elements, and the ICA regulon is a good supplement to the missing information. With better understanding of regulatory interactions, more useful features like the DR motifs of ArcA can be extracted from the sequence to explain regulons and improve poor performing machine learning models.

**Figure 3.**
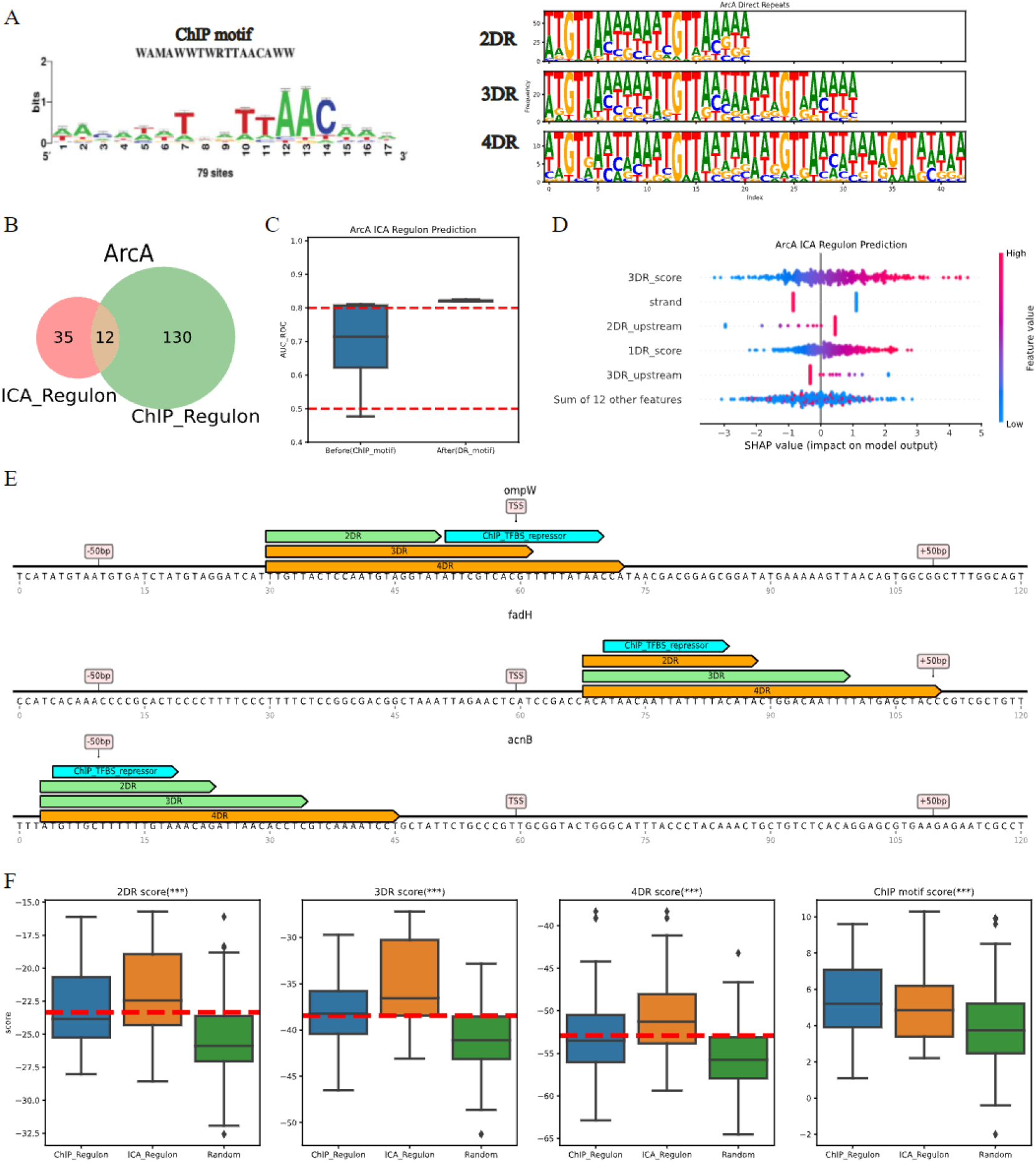
ArcA case study demonstrates how regulator-specific features can improve model performance. (A) ChIP motif of ArcA TF binding sites versus multiple direct repeats (DR) motifs. (B) The venn diagram of ArcA ChIP and ICA regulons. (C) Comparison of prediction accuracy with original feature matrix and multiple DR features. (D) Feature importance of ArcA ICA regulon prediction assessed by SHAP values. (E) Multiple DR motifs aligning to promoter sequences. Green: motif score above cutoff, Orange: motif score below cutoff, Blue: experimentally determined binding sites. Cutoff values are average motif scores of confirmed DR motifs. (F) Distributions of multiple DR motif scores and ChIP motif scores for ICA/ChIP regulons and random sequences.

### Understanding complex regulons through model variable importances

Five separate ICA-determined regulons are associated with the transcription factor Fnr. In a previous study, These have been named Fnr+IHF+gcvB, Fnr+NarL and Fnr+NarLP, Fnr-1 and Fnr-2, based on enrichment of different regulons within the ICA-determined regulon (12). Compared to Fnr+IHF+gcvB, Fnr+NarL and Fnr+NarLP, Fnr-1 and 2 contain much noise, which might be a factor leading to low prediction accuracy (**Figure 4AB**). IHF shape features (transformed by LDA) and Lrp motif score are two important features to determine Fnr+IHF+gcvB; for Fnr+NarL, important features are NarL and Fnr-1 IM motif scores (Fnr-1 IM motif resembles Fnr ChIP motif, **see S4 in SI**); and for Fnr+NarLP, they are NarL,NarP and Fnr-1 IM motif scores (**Figure 4C**). The distributions of features validate feature importances assessed. Fnr motif scores in ICA regulons have higher distributions than random sequences; Fnr+NarLP has both the highest NarL and NarP motif scores, Fnr+NarL has almost as high NarL motif scores but relatively lower NarP motif scores; And Fnr+IHF+gcvB’s IHF DNA shape feature has a significantly distinguishing distribution (**see SI, S5**).

**Figure 4.**
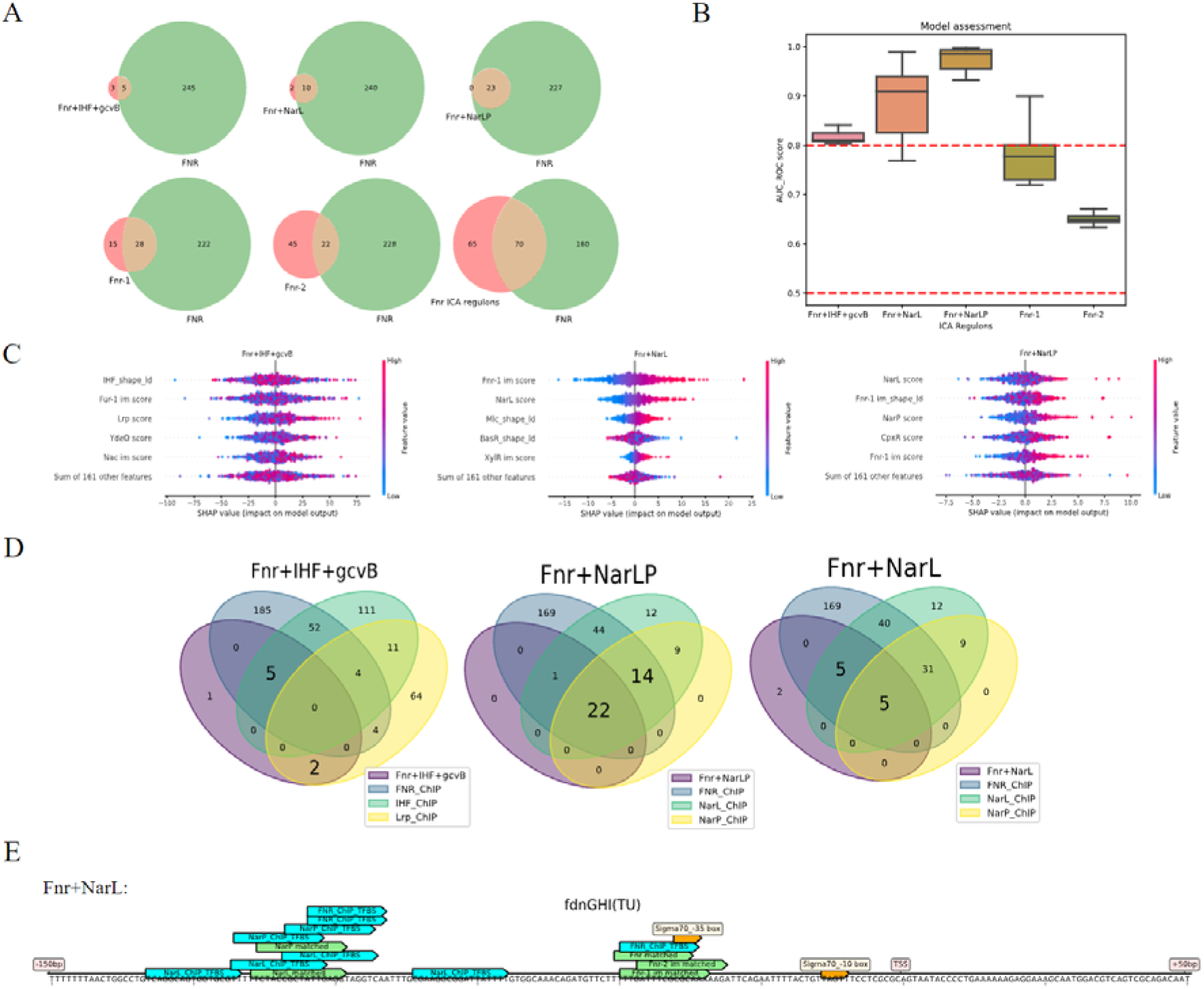
Fnr case study demonstrates the decomposition of complex regulons with machine learning. (A) Venn diagrams showing Fnr enrichment in 5 Fnr associated ICA regulons. (B) Assessment of predictive models for 5 Fnr associated ICA regulons. (C) Feature importances of predictive models for Fnr+IHF+gcvB, Fnr+NarL and Fnr+NarLP. (D) Venn diagrams of Fnr+IHF+gcvB, Fnr+NarL and Fnr+NarLP and ChIP regulons of Fnr,NarL,NarP,IHF and Lrp. (E) Promoter sequences of transcription units in Fnr associated ICA regulons.

Genes in Fnr+NarLP are mostly coregulated by Fnr, NarL and NarP, while only half of genes in Fnr+NarL are coregulated by Fnr,NarL and NarP, and the other half are just coregulated by Fnr and NarL; Fnr+IHF+gcvB mainly contains genes regulated by Lrp and coregulated by Fnr and IHF (**Figure 4D**). Looking at genes coregulated by the three TFs (Fnr, NarL and NarP), there are 14 genes not included in the Fnr+NarLP ICA regulon. We found that significantly higher NarL and NarP motif scores of genes in Fnr+NarLP appear to be a driving factor in this separation (**see SI, S6**). Furthermore, from feature positions on the sequence (**Figure 4E**), it could be observed that overlapping regulator motifs appear to be an underlying factor for the overlap in the observed regulons. This suggests that the machine learning model is able to summarize complex sequence features from multiple regulators into a prediction of regulon membership.

### Comparison between TRNs from ChIP experiment and ICA

Previous case studies of ArcA and Fnr observed that the distribution of motif scores tends to be higher in ICA regulons than in ChIP regulons. In order to validate this observation, 6 single TF-associated ICA regulons were compared to ChIP regulons. Motif scores of genes in the overlapping region of ChIP and ICA regulons are significantly higher than those of the remaining genes in ChIP regulons. In certain cases, such as Cra and CysB, the non-overlapping ICA regulon also has a higher motif score distribution than that of the non-overlapping ChIP regulon (**Figure 5B**). Thus, ICA regulons tend to extract genes with stronger binding sites. Furthermore, ICA regulons may contain regulator binding sites with high motif scores that remain undetected by experiments.

**Figure 5.**
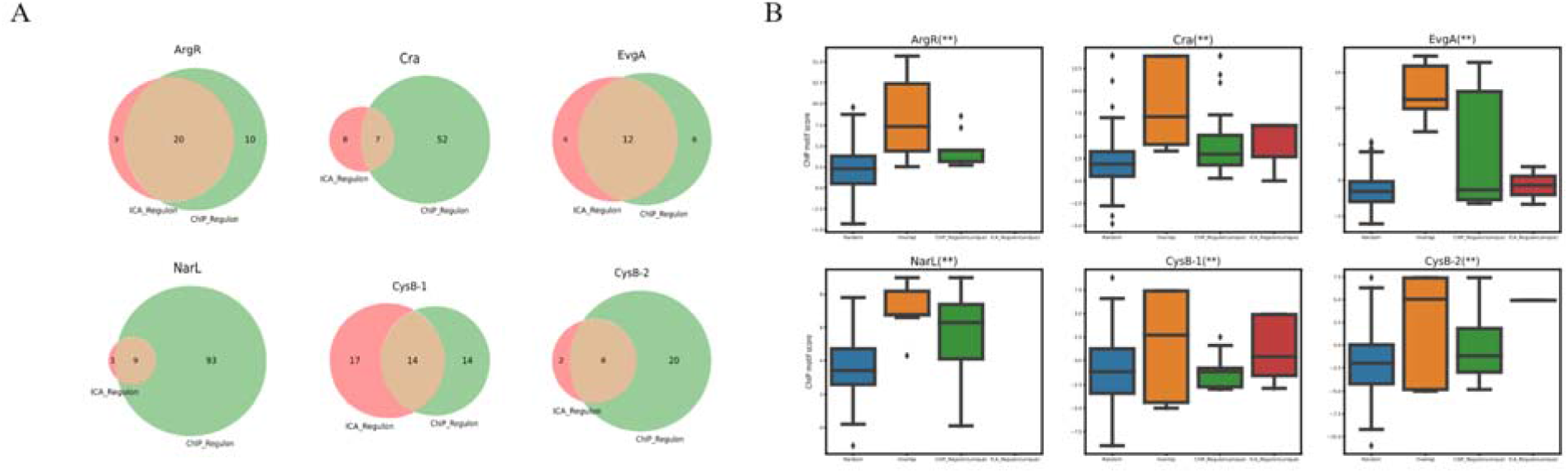
Comparison between ICA and ChIP-estimated regulons. (A) Venn diagrams showing consensus (brown) and unique (red/green) regulons estimated by each method. (B) Comparison of ChIP motif scores in the consensus regulon and unique regulons. Genes in the consensus regulons display a markedly higher distribution of motif scores.

### Sequence basis of TRN variation across strains

To explore TRN variation across multiple *E. coli* strains, we reconstructed the Fur pan-regulon of genes and transcription units for 6 strains across 3 phylogroups (**Figure 6AB**). The S/N ratios of Fur binding peaks from ChiP experiments in the core regulon were compared. Strains W and KO11FL in the F phylogeny group have higher Fur binding peak S/N ratios compared to MG1655, while CFT073 in B2 group has relatively low S/N ratios, indicating lower regulatory activity (**see SI, S8**). Strains in the same phylogroup tend to have similar regulatory activity.

**Figure 6.**
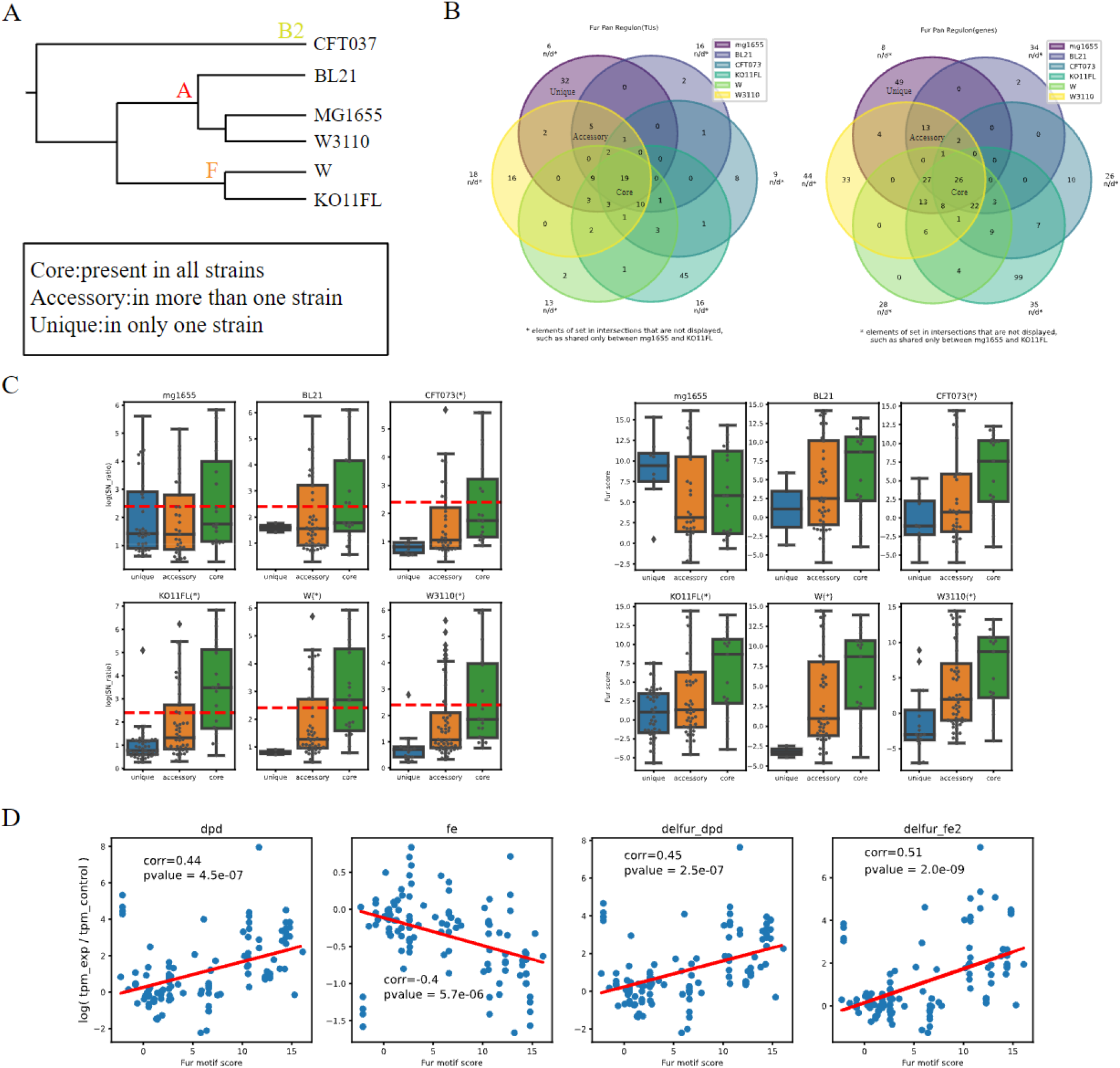
Fur pan-regulon analysis. (A) The phylogenetic tree of 6 strains investigated. (B) Fur pan regulons for both regulated genes and transcription units for 6 strains, with the phylogenetic tree displayed and unique,accessory, core regulons annotated. (C) Comparison of S/N ratios and Fur motif score: core > accessory > unique. (D) Expression change(————) versus Fur motif score. Regulatory response correlates well with the motif score.

Genes with ChIP-exo peaks in promoters are grouped into those with high and low S/N ratios at a cutoff of 10 (**see SI, S9**). While genes of low S/N ratios overlap little with Fur ChIP/ICA regulons, genes of high S/N ratios have large overlap with both types of regulons (**see SI, S10**). In addition, Fur ICA regulons determined with MG1655’s RNA-seq expression data appear robust at capturing genes of high S/N ratios across multiple strains (**see SI, S11**), which is consistent with the finding in the previous section that the ICA-based TRN contains genes of strong regulatory activity.

In the Fur pan-regulon, only genes in the core regulon are regulated in each strain; unique and accessory regulons vary across strains. Fur regulatory activity tends to be highest for core regulon genes, followed by the accessory regulon, and finally by the unique regulon. Motif scores follow a similar trend. These results suggest that binding site strength is the major factor causing regulatory activity variation between the core, accessory, and unique regulons (**Figure 6C**). Conserved promoters across strains tend to have high binding site strength, and thus high regulatory activity. Furthermore, a higher motif score indicates a stronger regulatory response, characterized by the expression change between control and experiment conditions (**Figure 6D**). These tendencies further validate that motif score is a good indicator of regulatory activity.

When the Fur pan-regulon is filtered just for genes in the high S/N ratio group, the conservation of high S/N ratio genes in each strain is not high enough to ignore TRN variation across strains, as the core region is usually below 30% (**Figure 7BC**). Therefore, a predictive model based on promoter features would be required to determine regulons of high S/N ratios in multiple strains. Trained with MG1655 strain’s feature matrix and target label indicating high and low S/N ratios, an LR classifier takes in other strain’s feature matrices and makes predictions (**Figure 7A**).

**Figure 7.**
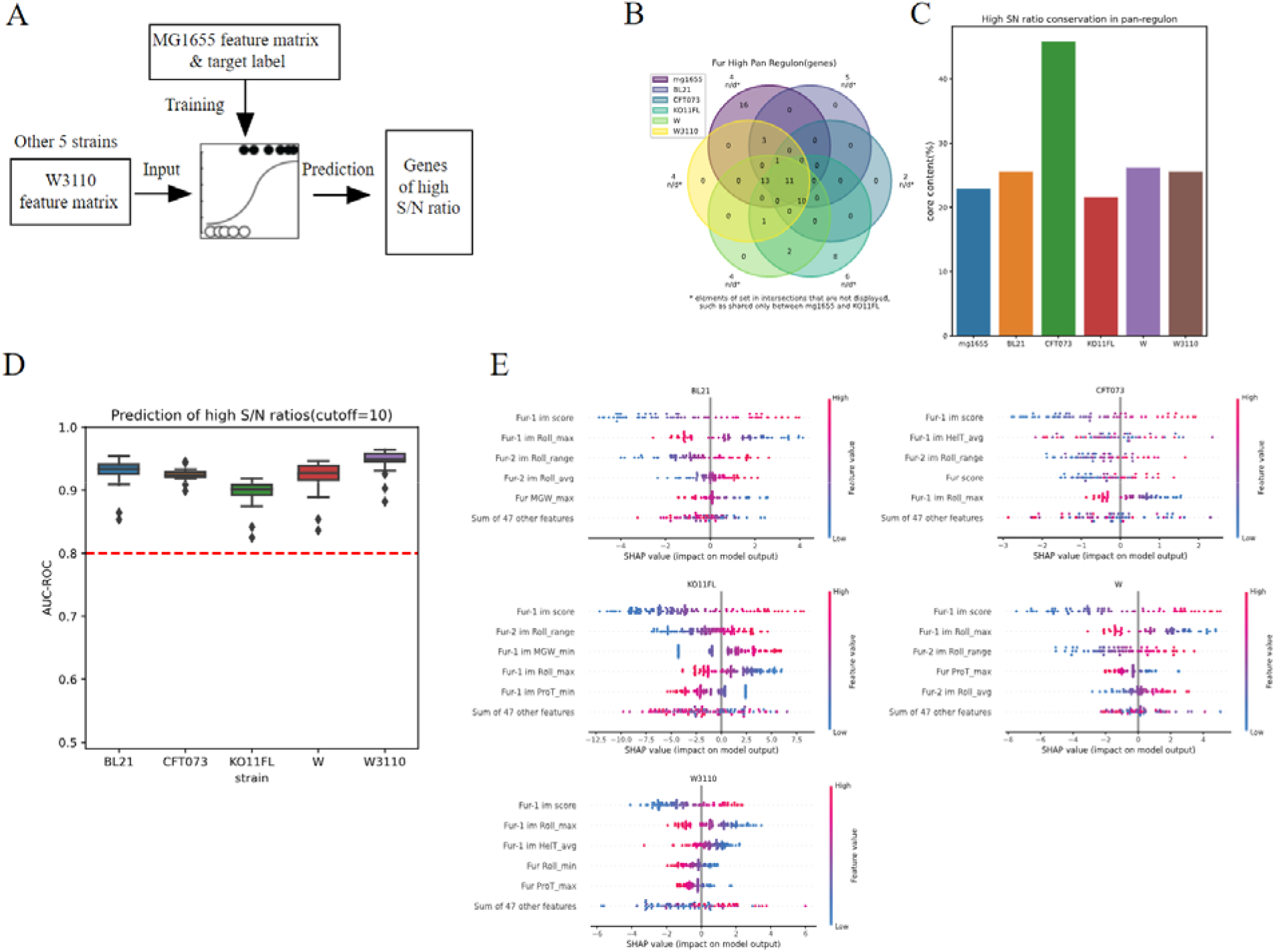
The prediction of multi-strain regulons: high and low S/N ratios. (A) The workflow of the predictive model.(B) Fur pan-regulon filtered for genes of high S/N ratios (C) Conservation of high S/N ratio genes in each strain. (D) Assessment of predictive models for 5 strains. (E) Feature importances: Motif scores and Roll are two most important features to predict high or low S/N ratios.

The prediction result demonstrates that the promoter sequence-based features used previously are sufficient to classify S/N ratios being high or low (**Figure 7D**). High correlation between motif scores and S/N ratios suggests that motif scores are important indicators (**see SI, S12**). The range of DNA roll degree is another important indicator, as it appears in all models’ most important features (**Figure 7E**). The distributions of DNA roll degree range at matched binding sites between genes of high and low S/N ratios show the tendency that gene promoters’ binding sites with larger roll degree range are more likely to have high S/N ratios (**see SI, S13**).

In summary, the sequence basis of TRN’s variation across strains lies in binding site strength, characterized by motif scores. Also, the important features for classifying S/N ratios as high or low are robust across strains: motif scores and DNA roll degree range.

## Discussion

We developed DNA sequence-based machine learning models for gene regulon membership, resulting in high prediction performance for 57/68 of possible *E. coli* ICA-determined regulons. Successful models highlight that gene transcriptional regulation can be quantitatively explained by relatively simple promoter sequence features for many regulons. We found that TF motif scoring and DNA shape are both critical for determination of top-down or bottom-up regulon membership. However, certain regulons require additional specialized transcription factor-specific sequence features, exemplified by the need to add ArcA direct repeat motifs to obtain a good model for the ArcA regulon.

Generally, a higher motif score indicates higher similarity to the consensus sequence, but it only accounts for binding site strength with positional frequency, leaving out other properties of DNA sequences. Therefore, the motif score is important but not sufficient to determine regulon membership. To characterize regulator-sequence interactions structurally, DNA shape features were included in the feature matrix for machine learning models, and they acted as significant features in the prediction, consistent with previous work in this area (11). TF-specific DNA shape vectors reflect unique regulator protein structures and provide another measurement for regulator binding affinity.

Comparing these two types of regulons demonstrates that ICA regulons identify genes of strong binding site strength. If a gene is included by both types of regulons, the regulatory interaction is not only assessed by ChIP based experiments but also validated by unraveled sources of transcription signals. Genes unique to ICA regulons also have high motif scores, indicating potentially undetected parts of the TRN. Similarly, binding site strength and S/N ratios also distribute unevenly across unique, accessory and core regions in the Fur pan-regulon: the core regulon is more active than the unique regulon due to high conservation level of promoters in core regulon. Considering the relatively larger overlap between high S/N ratio sites and two types of regulons,the S/N ratio might also be an indicator of the confidence level, in which case, the regulatory interactions in core regulon have a higher confidence level.

In the analysis of the Fur pan-regulon, promoters in other strains of *E. coli* must be predicted, but the method used in this study could only find promoters in highly conserved transcription units, and hence, many promoters’ positions are unknown. The lack of promoter data resulted in the infeasibility of computing sequence features for certain genes, which led to filtration of measured S/N ratios. The final result was also affected: many genes in MG1655’s unique regulon of low S/N ratios were filtered out due to missing transcription start sites. Therefore, a novel method to predict promoters is needed. If the previously mentioned scoring function using both PSSM and DNA shape is applicable, it can predict −10/−35 boxes of sigma factors with high accuracy. Since the distance between those 2 boxes to the transcription start site is stable, the promoter could also be determined.

There is substantial room for improvement of model performance for poorly performing regulons. Models for 32 ICA regulons could not be determined due to lack of necessary information for feature computation, such as regulator binding site motifs for small regulons or transcription start site annotation. Improvements on sequence feature computation will improve the accuracy of machine learning and thus more top-down regulons would be explainable. A better motif matching method than linear search with PSSM could be developed to improve the computation of motif scores and better characterize binding site strength. Though PSSM provides the probability of the segment being a binding site, the shape of the DNA sequence is not considered. There have already been a few studies using DNA shape in binding site prediction, for example, improvement was observed in the machine learning model using both DNA sequence and shape to predict TF binding sites in ChIP-seq datasets (14). Therefore, predicting a TRN will become more accurate if the scoring function used in motif matching can be formulated with both PSSM and DNA shape information. In addition, further characterization of TF-specific sequence features could improve prediction of regulons with poorly performing models. For instance, multiple direct repeats motifs of ArcA characterize diverse binding site architecture better than the ArcA original ChIP-determined motif, and inclusion of this ArcA-specific sequence feature improves the prediction accuracy for the ArcA ICA regulon. Given enough data, the machine learning workflow should be able to reconstruct the TRN with high accuracy.

Taken together, this study introduces a reliable workflow to analyze transcriptional regulation activity in prokaryotes based on promoter sequence, using machine learning models to reconstruct the TRN. Biophysical factors affecting TRN architecture and regulatory activity were surveyed, offering directions for data-driven genome design in synthetic biology to control cellular activity by tuning sequence features.

## Methods

### The Bitome: Curating and organizing genomic feature information

Genome information for *Escherichia coli* K-12 MG1655 was collected and curated from the NCBI RefSeq and RegulonDB databases. The Bitome Python software package was utilized to load and organize sequence-based information from these data sources, including genome sequence, coding sequences, transcription start sites, transcription units, and transcription factor binding sites (15). The code containing workflows used in this work is available at https://github.com/SizheQiu/IM-ML.

### Motif score of promoter sequence

A position-specific scoring matrix, or PSSM, was generated for a given TF by aligning its known binding site sequences and computing the relative probability of finding each DNA base at each binding site position. As the PSSM’s coefficients are the log-odds of a probability, the summation of the log-odds at all nucleotide positions in a putative binding site represents the probability of motif existence. The formula is 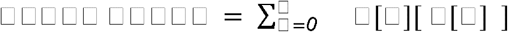, and M is PSSM, k is motif length, S is the segment, i is the index. The algorithm performs 1-D dynamic programming that iteratively computes the motif score of consecutive equal-length segments (length = motif length), and the segment of highest score is the matched motif box. Then, the highest score is output as the motif score of the promoter sequence.

For transcription factors, the search range was −150bp and +50bp to the transcription start site, and in this study, PSSMs were obtained from RegulonDB’s TF PWMs Browser page, mainly determined experimentally by ChIP seq experiments (16). Unlike TF motifs, sigma factor motifs consist of two binding regions with a gap of varying length: the −10 box (Pribnow box) and −35 box. PSSMs of −10 and −35 boxes for sigma 70/32/38/24/54/28 were generated from annotated sigma factor binding sites in RegulonDB and consensus sequences from the reference textbook *Transcription regulation in prokaryotes (17)*. Other sigma factors were not included due to limited annotation. The search range was 20bp upstream for −10 box, 40bp for - 35 box. In addition to motif scores of −10/−35 boxes on promoter sequences, an AT-rich 7bp segment in the spacer between −10 and −35 boxes was matched and the highest AT ratio was used to account for the extended −10 box (18). Sigma factor related features were motif scores and Hamming distances of −10/−35 boxes, the distance from −10/−35 boxes to the transcription start site, spacer length, AT ratio of the spacer, and AT ratio of the extended −10 box. To reduce feature dimensions, a hyperplane decision surface, dividing the sigmulon (genes regulated by the sigma factor) and other genes, was computed using LDA (linear discriminant analysis), and features were projected to it.

Some motifs in RegulonDB failed to capture the whole binding site consensus sequence, or other motifs also important to regulation activity were missed. For example, the Fur binding site motif in RegulonDB didn’t include the whole 19bp length inverted complementary repeats structure (19). Motif discovery in ICA regulons (iModulons) using MEME suite provided motifs missed by RegulonDB, named as “ IM motifs “in contrast to “ ChIP motifs “. MEME is a toolbox used to discover novel motifs in collections of unaligned sequences (20). Those ICA regulon motifs were used to supplement information missed by ChIP experiments.

Another possible edge case was that the regulator binding site’s architecture has great diversity, which could not be characterized by a simple motif. For instance, 42% of ArcA binding sites are 2 direct repeats of 10bp segments, 41% are 3 direct repeats, 15% are 4 direct repeats and 5% are 5 direct repeats (13). And hence, direct repeats (DR) motifs were used to compute features for model performance improvement.

### DNA shape feature computation

In this study, 13 types of shape features were computed using DNAShapeR, a DNA shape predictor in R language (21). These shape features included 6 inter-base pair shapes: Roll, HelT (Helix Twist), Shift, Slide, Rise and Tilt, and 7 intra-base pair shapes: MGW (Minor Groove Width), ProT (Propeller Twist), Buckle, Shear, Stretch, Stagger, Buckle and Opening.

A pentamer query table integrated with a sliding-pentamer window was used to compute the shape vectors for all matched TF binding sites. The pentamer table was obtained from the GitHub repository of DNAShapeR. For predicting intra-base pair parameters, each sliding step assigned a shape prediction for the central base pair. For predicting inter-base pair parameters, each sliding step assigned a shape prediction for two central base-pair steps. The overlapping values arising from two adjacent pentamers at the same nucleotide position were averaged. The sliding-window approach thus results in a shape vector (22). Maximum, minimum, range, and mean values of the shape vector were computed as features. To reduce dimensions of shape features, LDA was applied to project shape features onto a hyperplane decision surface, which divides the ChIP regulon and other genes.

### Logistic regression classification on ICA regulon membership and assessment of model performance

A logistic regression classifier was chosen in this study because it is a more interpretable model compared to common alternatives such as Random Forest or Neural Network models, and the degree of overfitting is usually smaller. The classifier was implemented using the scikit-learn Python package (23). The target labels were binarized ICA regulon memberships. Two feature matrices were trialed: only ChIP motif scores, and all features. Model performance was assessed by 5-fold cross validation and area under the curve of receiver operating characteristic (AUC ROC), showing true positive rate versus false positive rate across all possible classification thresholds. A cutoff of 0.8 was used to indicate good performance. For optimal classification performance, hyperparameters of the LR classifier were randomly searched: l1 or l2 penalty, C parameter(inverse of regularization strength) ranging from 0.1 to 100, and maximum iterations ranging from 100 to 2000. The combination of hyperparameters with the highest AUC ROC would be selected.

### Promoter prediction and pan-regulon reconstruction

Though genomic information of *E. coli* MG1655 is well annotated, other strains, such as W3110, have limited genomic objects annotated, especially the lack of transcription start sites that indicate the location of promoters. Therefore, predicting promoter locations was necessary for the multi-strain study of *E. coli.* We aligned sequences of transcription units in MG1655 to other strains’ genome sequences using BLAST (24). The other strains studied were: KO11FL, CFT017, W, W3110, and BL21. Inexact alignments were filtered out; only exact matches were used to map MG1655 transcription units onto the genomes of other strains. In this manner, transcription start sites were also determined for all strains.

After promoters were predicted for each strain, Fur ChIP-exo S/N ratios were mapped to promoter sequences, and transcription units regulated by Fur in each strain were determined. Since predicted transcription units were all exact matches, the genes in transcription units were known, therefore genes regulated by Fur were also determined. The Fur pan-regulons of genes and transcription units was then reconstructed for the six strains of *E. coli:* MG1655, KO11FL, CFT017, W, W3110 and BL21, across three phylogroups: A, B2 and F. Unique, accessory and core regulons were annotated based on shared Fur regulation across strains.

### Logistic regression classification on S/N ratios high or low of Fur ChIP-exo sites in multiple strains

The LR classifier was implemented in the same way as in section 2.4, and performance was also assessed using AUC ROC. A cutoff = 10 (2.3 in log-scale) separated each strain’s S/N ratios for their bimodal distribution of Fur ChIP-exo sites into high and low values, which was binarized into the target label for classification. The resulting feature matrix included all features computed in section 2.2 and 2.3, the same as LR classification of ICA regulon membership. The *E. coli* K-12 MG1655 feature matrix and target label were used as the training set to build the predictive model for other strains’ S/N ratios being high or low. Hyperparameter optimization was done in the same manner as in section 2.4.

## Acknowledgements

We would like to thank Ye Gao for helpful discussions on this manuscript. This work was supported by the Novo Nordisk Foundation grant numbers NNF10CC1016517 and NNF20CC0035580.

## Notes

### Competing Interest Statement

The authors have declared no competing interest.

