## Supplementary Information for "Quantitative sequence basis for the *E. coli* transcriptional regulatory network"

**This PDF file includes:**

Figures S1 to S13
SI References

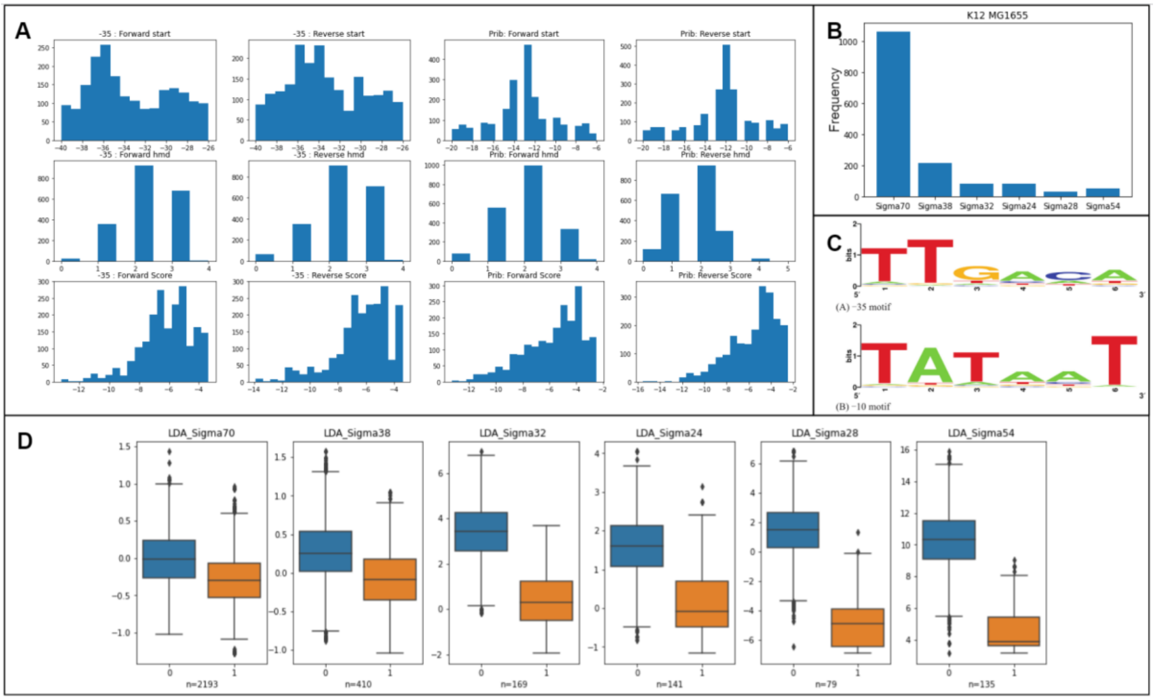


**Fig. S1.** Sigma factor binding boxes matching results. (A) Locations, hamming distances and motif scores of sigma factor 70's matched pribnow(-10) box and -35 box for . Consistent results for genes in reverse and forward strands. (B) The sizes of experimentally determined sigmulons: sigma factor 70/38/32/24/28/54. (C) Motifs of sigma factor 70's -10 and -35 boxes. (D) Distribution of sigma factor related features processed by LDA. 1 for genes in the sigmulon and 0 for other genes. Features include distances of -10/-35 boxes to transcription start sites, motif scores,hamming distances, spacer length, AT content of extended -10 box in the spacer.

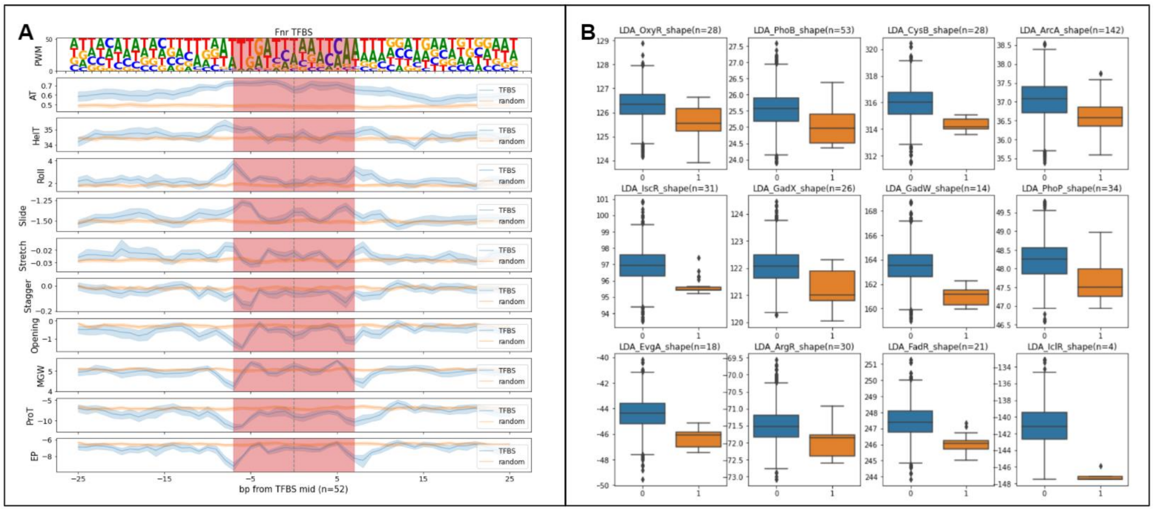


**Fig. S2.** DNA shape profile. (A) DNA shapes at Fnr TFBS. Shapes include HelT, MGW, ProT, roll, slide, stretch, stagger and opening. (B)Distribution of DNA shape features transformed by LDA. 1 for genes in ChIP regulon of the TF, and 0 for other genes.

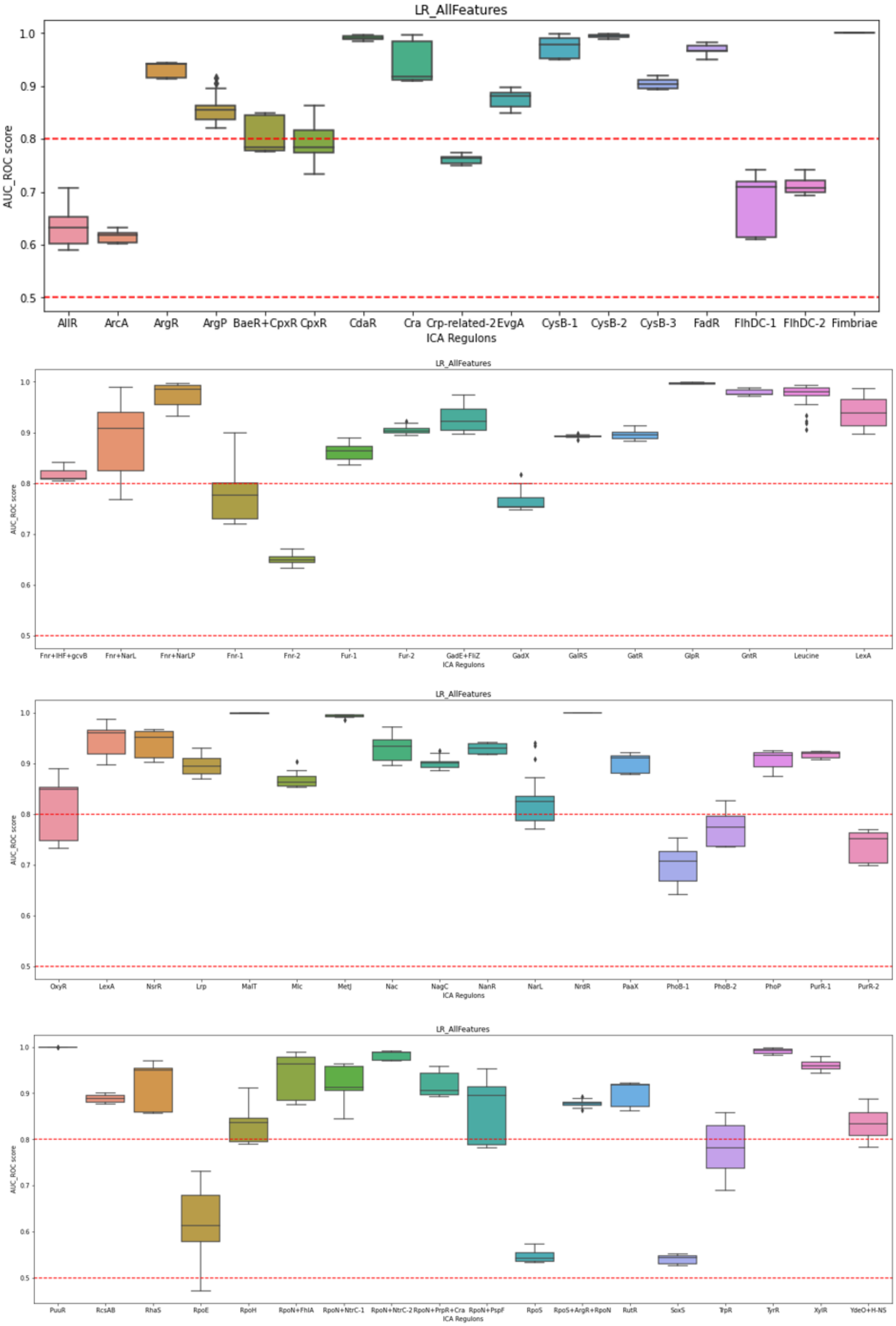


**Fig. S3.** Model assessment for 68 ICA regulons. The 0.5 AUC ROC is the threshold for baseline models, and 0.8 is threshold for good models.

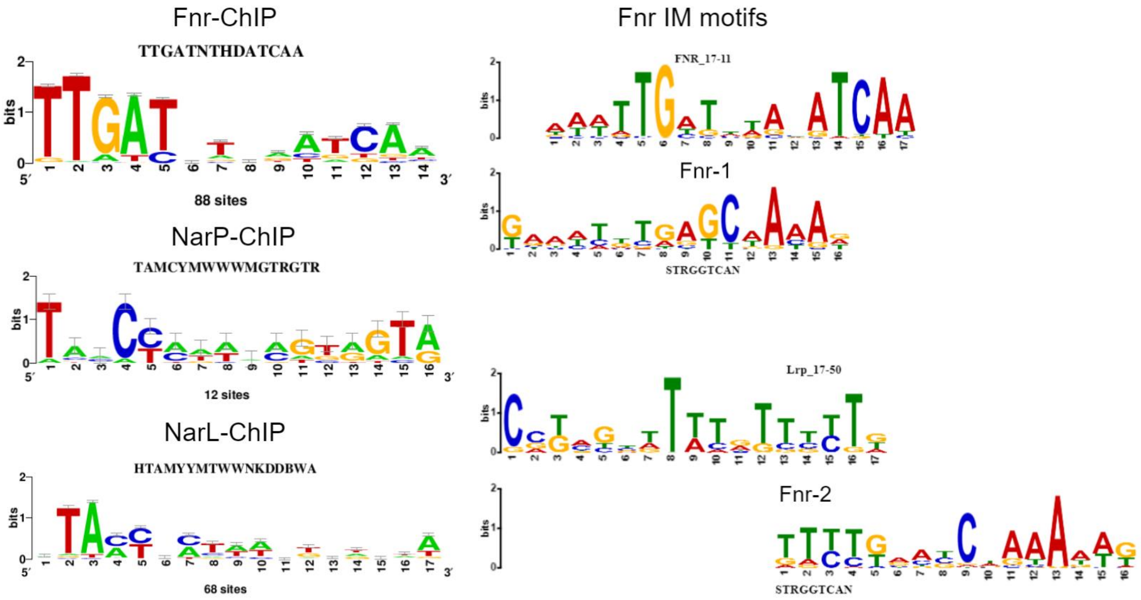


**Fig. S4.** Fnr,NarL and NarP ChIP motifs (left) and ICA regulon motifs of Fnr-1,2 (right). Most similar TFBS motifs for Fnr-1,2 IM motifs are displayed: Fnr-1 and Fnr-2 IM motifs are most similar to Fnr and Lrp TFBS motifs.

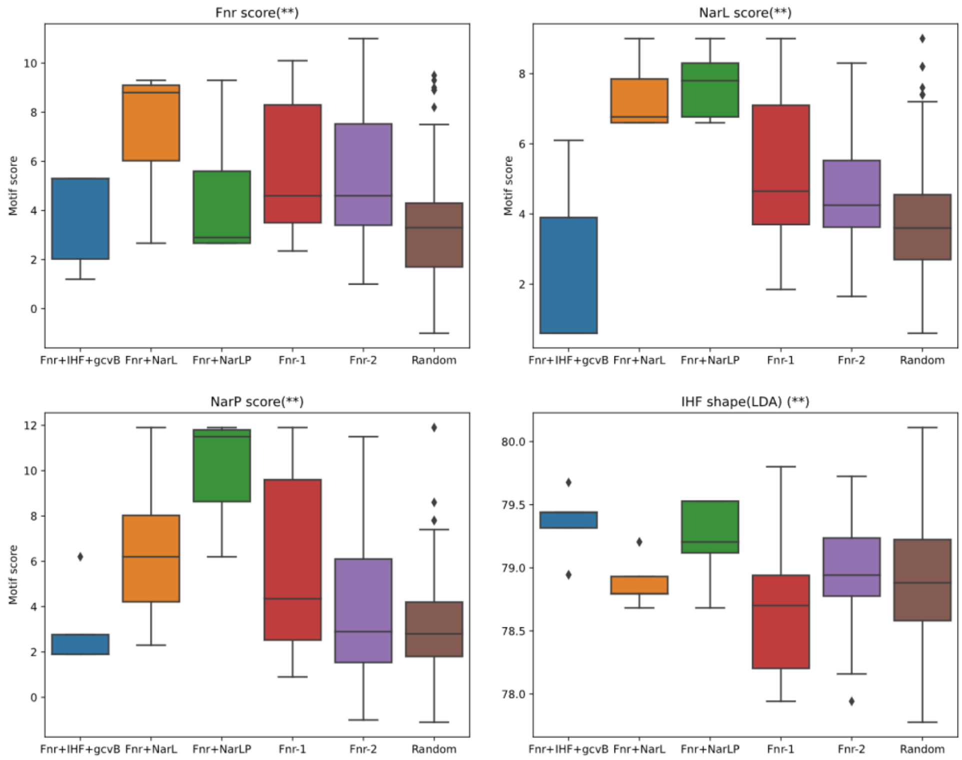


**Fig. S5.** Distributions of features across ICA regulons: Fnr, NarL and NarP motif scores and IHF shape features(transformed by LDA).

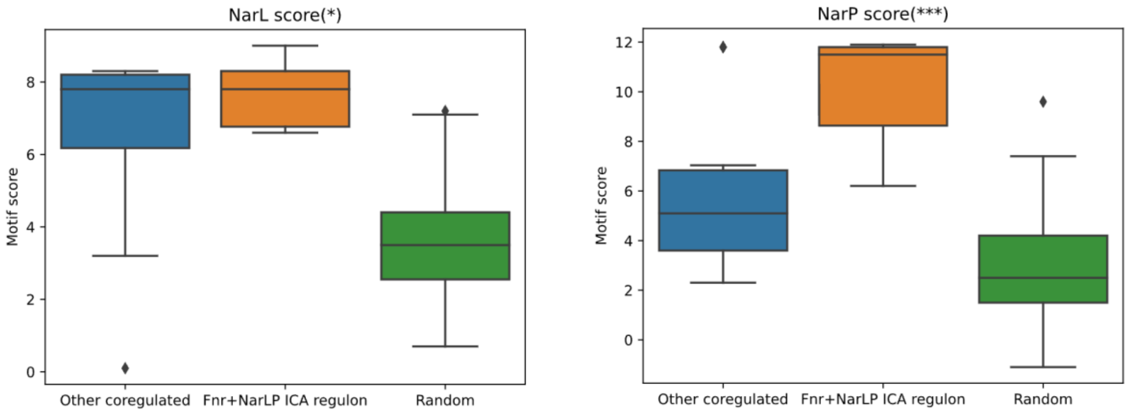


**Fig. S6.** Comparison of NarL and NarP motif scores between genes in Fnr+NarLP and other genes coregulated by Fnr,NarL and NarP.

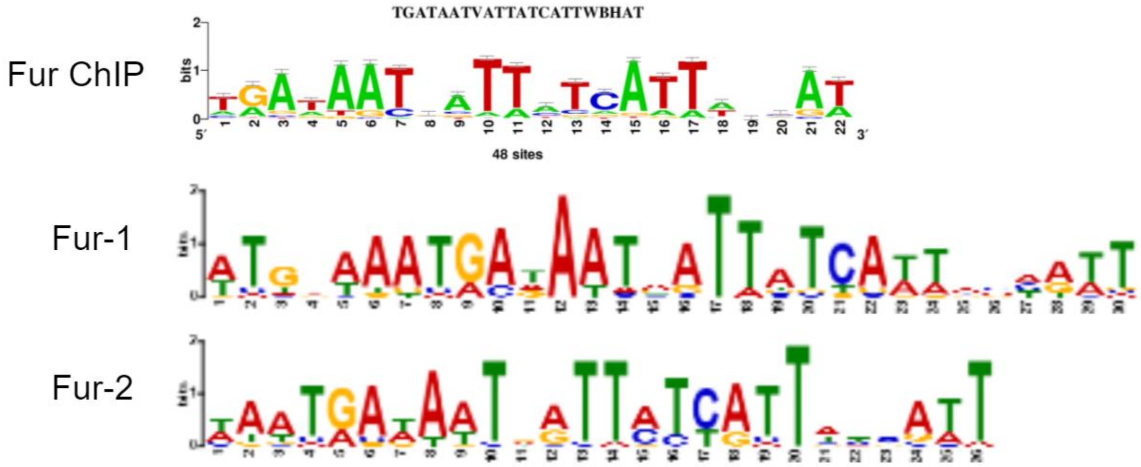


**Fig. S7.** ChIP and ICA regulon motifs of Fur in E.coli K-12 MG1655.

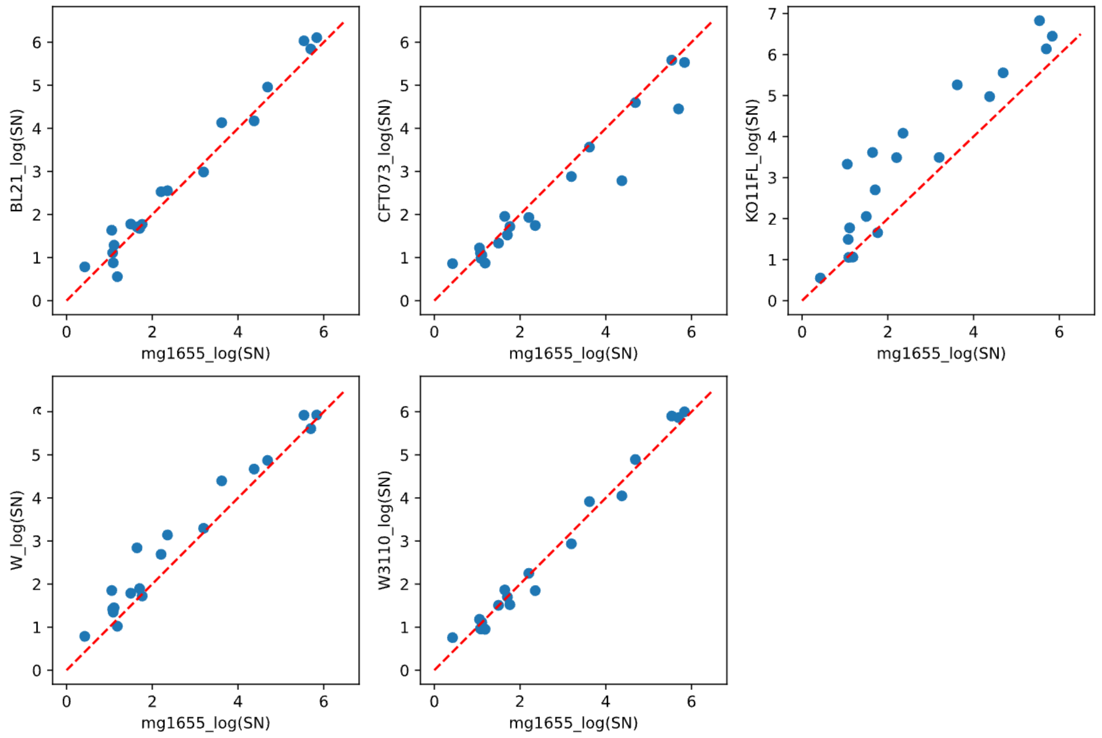


**Fig. S8.** Comparison of S/N ratios between MG1655 and all other 5 strains.

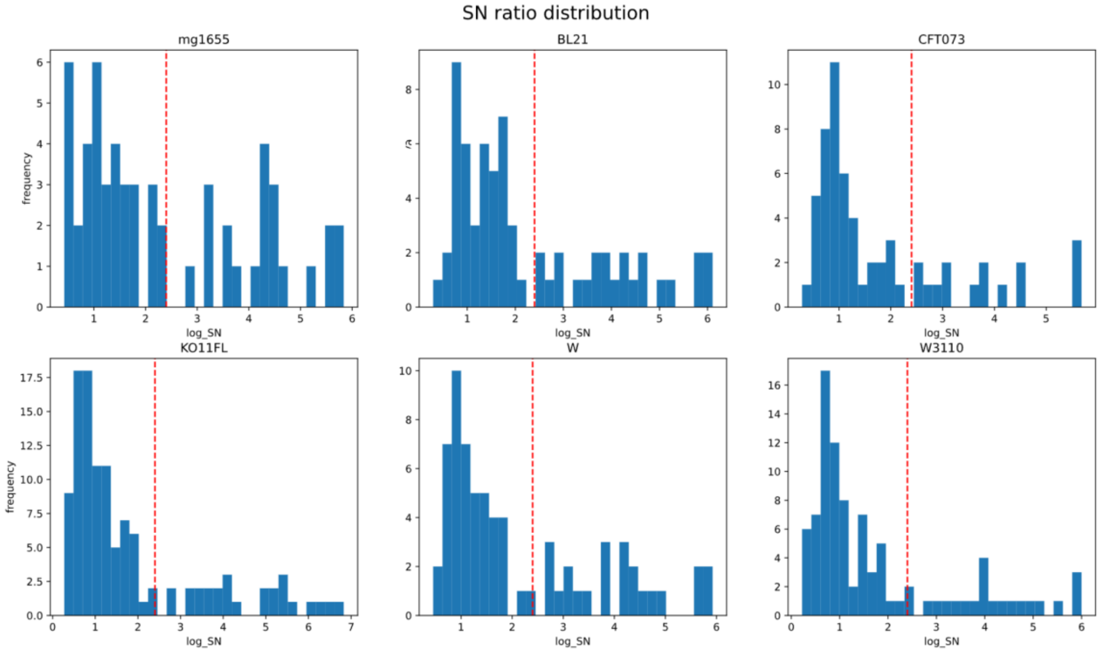


**Fig. S9.** Histograms of S/N ratios in each strain. Bimodal distribution can be observed and a cutoff = 10 (2.3 in log-scale) separates high and low S/N ratios.

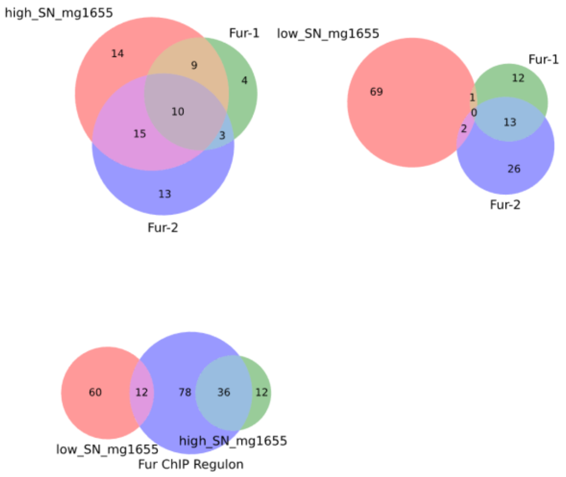


**Fig. S10.** Venn diagrams showing relationships of genes with high/low S/N ratios and Fur ICA/ChIP regulons for MG1655 strain.

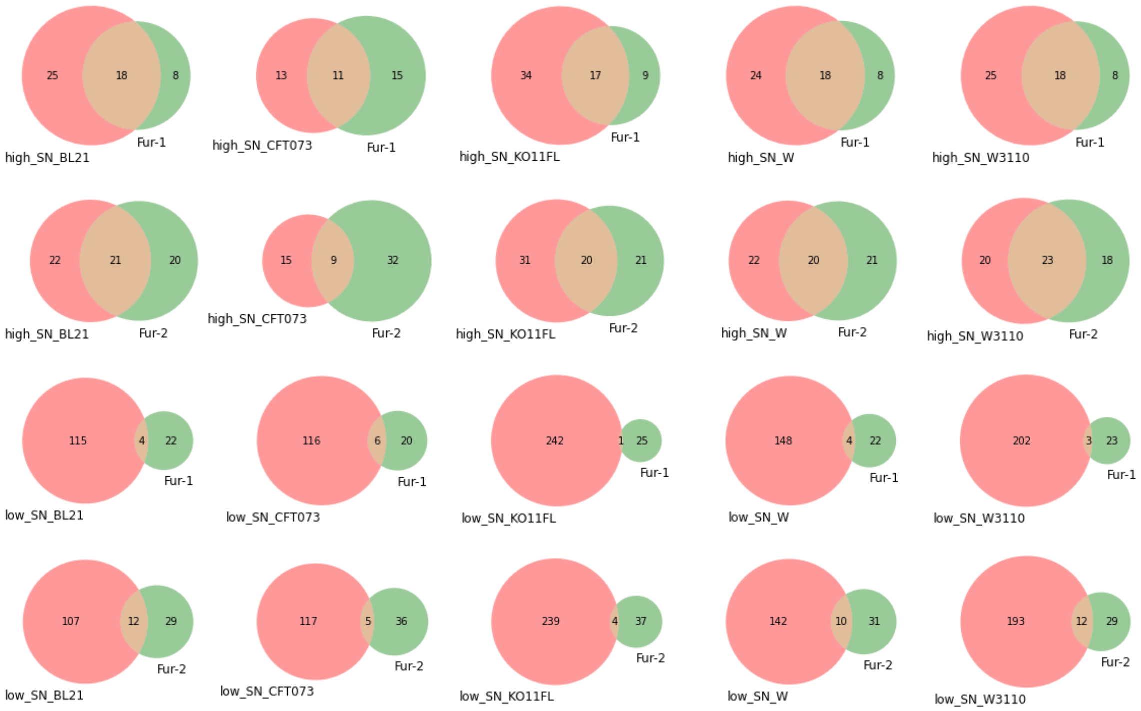


**Fig. S11.** Venn diagrams of Fur ICA regulons and Fur high/low SN ratio groups across multiple strains: BL21,CFT073,KO11FL,W and W3110.

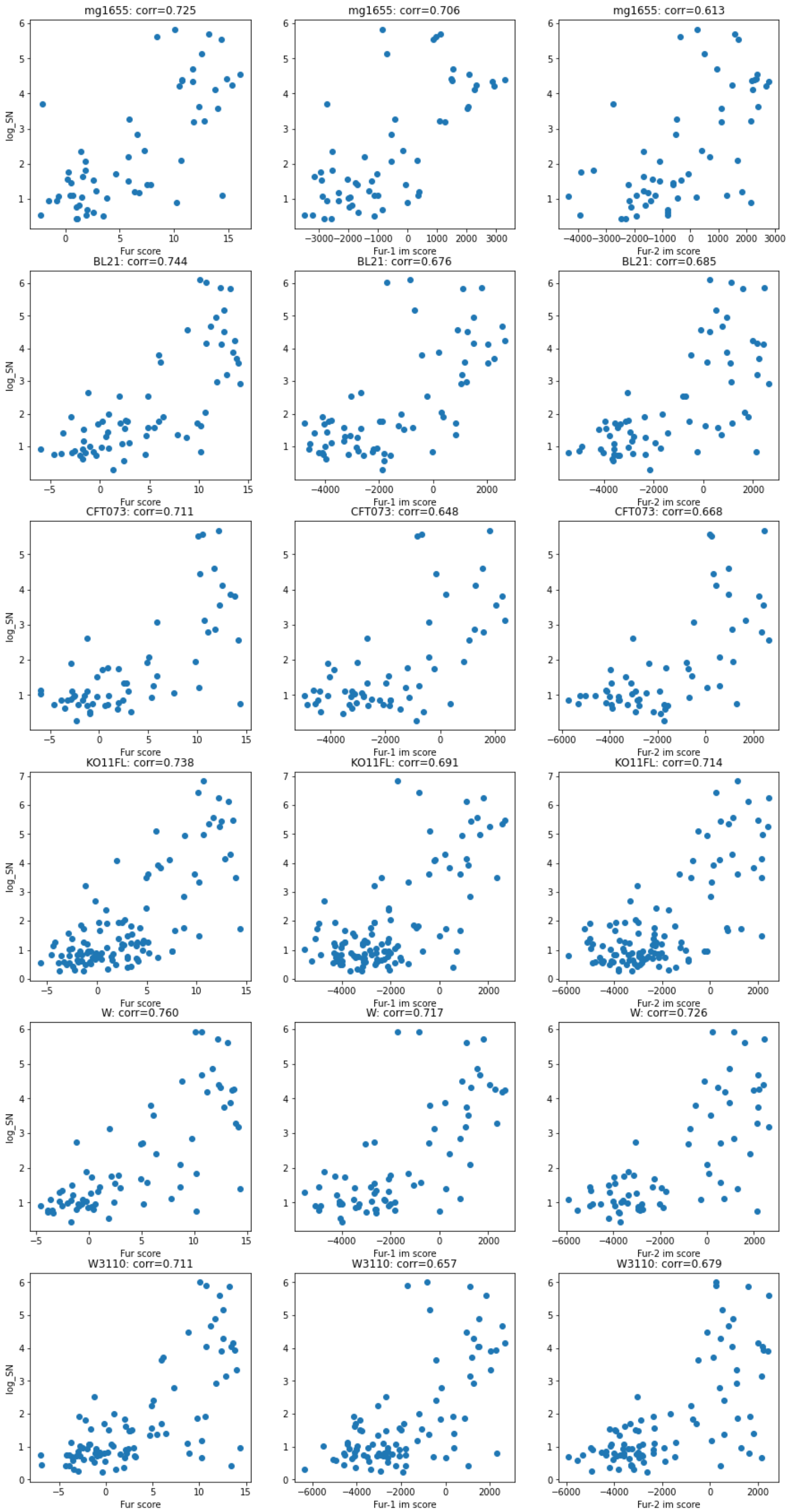


**Fig. S12.** Correlation of Fur motif scores (Fur ChIP motif and Fur-1,2 motifs) and S/N ratios across multiple strains: MG1655,BL21,CFT073, KO11FL,W and W3110

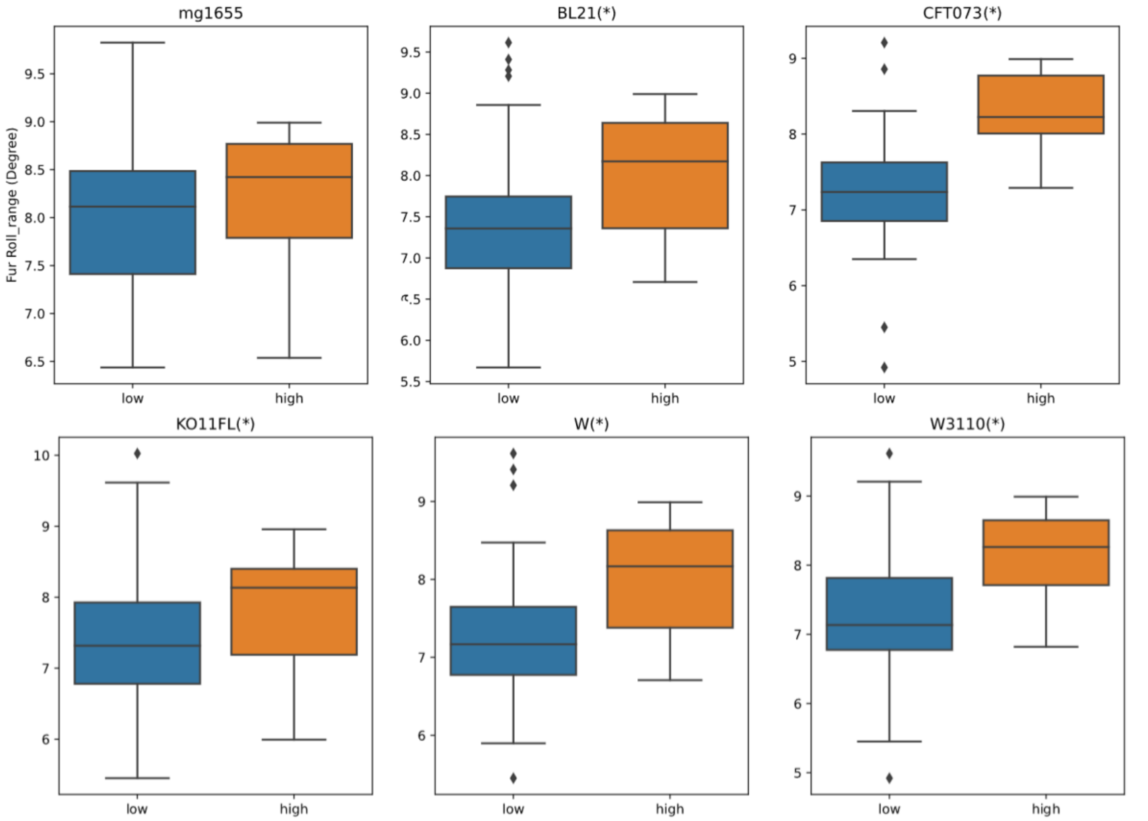


**Fig. S13.** Comparison of Roll range at matched binding sites between genes of high and low S/N ratios. The Larger Roll range indicates a higher S/N ratio.
